# Working memory enhancement using real-time phase-tuned transcranial alternating current stimulation

**DOI:** 10.1101/2024.05.31.596854

**Authors:** David Haslacher, Alessia Cavallo, Philipp Reber, Anna Kattein, Moritz Thiele, Khaled Nasr, Kimia Hashemi, Rodika Sokoliuk, Gregor Thut, Surjo R. Soekadar

## Abstract

**Background:** Prior work has shown that transcranial alternating current stimulation (tACS) of parietooccipital alpha oscillations (8 – 14 Hz) can modulate working memory (WM) performance as a function of the phase lag to endogenous oscillations. However, leveraging this effect using real-time phase-tuned tACS was not feasible so far due to stimulation artifacts.

**Objectives/Hypothesis:** We aimed to develop a system that tracks and adapts the phase lag between tACS and ongoing parietooccipital alpha oscillations in real-time. We hypothesized that such real-time phase-tuned tACS enhances working memory performance, depending on the phase lag.

**Methods:** We developed real-time phase-tuned closed-loop amplitude-modulated tACS (CLAM-tACS) targeting parietooccipital alpha oscillations. CLAM-tACS was applied at six different phase lags relative to ongoing alpha oscillations while participants (N = 21) performed a working memory task. To exclude that behavioral effects of CLAM-tACS were mediated by other factors such as sensory co-stimulation, a second group of participants (N = 25) received equivalent stimulation of the forehead.

**Results:** WM accuracy improved in a phase lag dependent manner (p < 0.05) in the group receiving parietooccipital stimulation, with the strongest enhancement observed at 330° phase lag between tACS and ongoing alpha oscillations (p < 0.01, d = 0.976). Moreover, across participants, modulation of frontoparietal alpha oscillations correlated both in amplitude (p < 0.05) and phase (p < 0.05) with the modulation of WM accuracy. No such effects were observed in the control group receiving frontal stimulation.

**Conclusions:** Our results demonstrate the feasibility and efficacy of real-time phase-tuned CLAM-tACS in modulating both brain activity and behavior, thereby paving the way for further investigation into brain-behavior relationships and the exploration of innovative therapeutic applications.

## Introduction

Working memory (WM) allows for the temporary storage and modification of information, such as remembering a phone number before writing it down. WM is therefore essential for many complex brain functions, such as arithmetic calculation [1], reading comprehension [2], spatial reasoning [3], learning [4], problem-solving [5], and goal-directed behavior [6]. Brain oscillations, particularly in the theta (4 – 8 Hz) and alpha (8 – 14 Hz) range, have been identified as a key mediator of WM accuracy [7–14]. Understanding the role of brain oscillations in WM is crucial to understanding how WM underpins cognition [15], and to developing interventions that restore cognitive functions in brain disorders involving WM impairments, such as dementia or attention deficit disorder [11, 16].

Transcranial alternating current stimulation (tACS) is a noninvasive neuromodulation technique that has been used to target brain oscillations in WM [11, 16–19] and improve WM function [11, 16, 20]. By applying sinusoidal electrical currents to the scalp, tACS can interact with neural oscillations in targeted brain regions [21–24]. Typically, tACS is applied at the frequency of endogenous brain oscillations thought to be entrained to the externally applied electric field [24]. While tACS has been used to demonstrate a causal link between theta oscillations and WM [17, 25], such causal link between WM accuracy and alpha oscillations remained elusive [26]. This may be due to the dynamic interaction between changing brain states and constant tACS waveforms [27–31]. For example, effects of tACS are known to depend on its phase lag to endogenous brain oscillations [29, 32–34], which continuously fluctuates throughout an experimental session. Since the phase of tACS waveforms is typically fixed, i.e., non-adaptive relative to the continuously fluctuating phase of the endogenous oscillation, interaction between the tACS waveform and endogenous brain oscillations may lead to both random enhancement and suppression of brain oscillatory activity and behavior [33]. Such a state-dependency of tACS is believed to be a major source of effect variability [27, 31, 33–35], limiting its scientific and clinical utility.

Recent work has demonstrated that the effects of tACS on parietooccipital alpha oscillations and WM accuracy depend on the phase lag between tACS and the ongoing oscillation [13]. In this study, brief trains (0.8 s) of tACS were triggered to align at either 0° or 180° phase lag to the ongoing alpha oscillation. It was found that WM accuracy was higher when tACS was triggered at 0° compared to 180° phase lag. Importantly, stimulation artifacts impeded the assessment of brain activity in simultaneously recorded electroencephalography (EEG), such that tACS could not be adapted to ongoing alpha oscillations in real-time. To leverage phase-tuned tACS for the modulation of brain activity and behavior, continuous adaption of the tACS waveform would be necessary, e.g., to maintain the phase lag between tACS and endogenous brain oscillations [33, 34, 36].

## Materials and methods

First, phase tuning of closed-loop amplitude-modulated transcranial alternating current stimulation (CLAM-tACS) to parietooccipital alpha oscillations was validated. Subsequently, the relationship between the CLAM-tACS phase lag and WM accuracy was assessed. Moreover, optimal phase to enhance WM accuracy was evaluated separately for each participant. In accordance with prior work [27], the link between modulations of long-range synchrony of alpha oscillations as measured by phase lag index (PLI) and modulations of WM accuracy was assessed.

### Experimental paradigm

For validation of real-time phase-tuned CLAM-tACS (Fig. 1), we used a visual working memory task. Assessment of brain oscillations during CLAM-tACS was achieved using stimulation artifact source separation [37]. CLAM-tACS was applied at six different phase lags to parietooccipital alpha oscillations while participants performed a visual WM task (Fig. 2A). To assess whether effects of CLAM-tACS (Fig. 2B) were mediated by sensory co-stimulation (e.g., retinal, or somatosensory stimulation), we implemented a control condition where all experimental parameters remained identical except that the stimulation electrodes were placed on the forehead (Fig. 2C). To quantify effects of CLAM-tACS, we assessed WM accuracy and long-range synchronization of alpha oscillations as measured by the PLI during stimulation.

**Figure 1.**
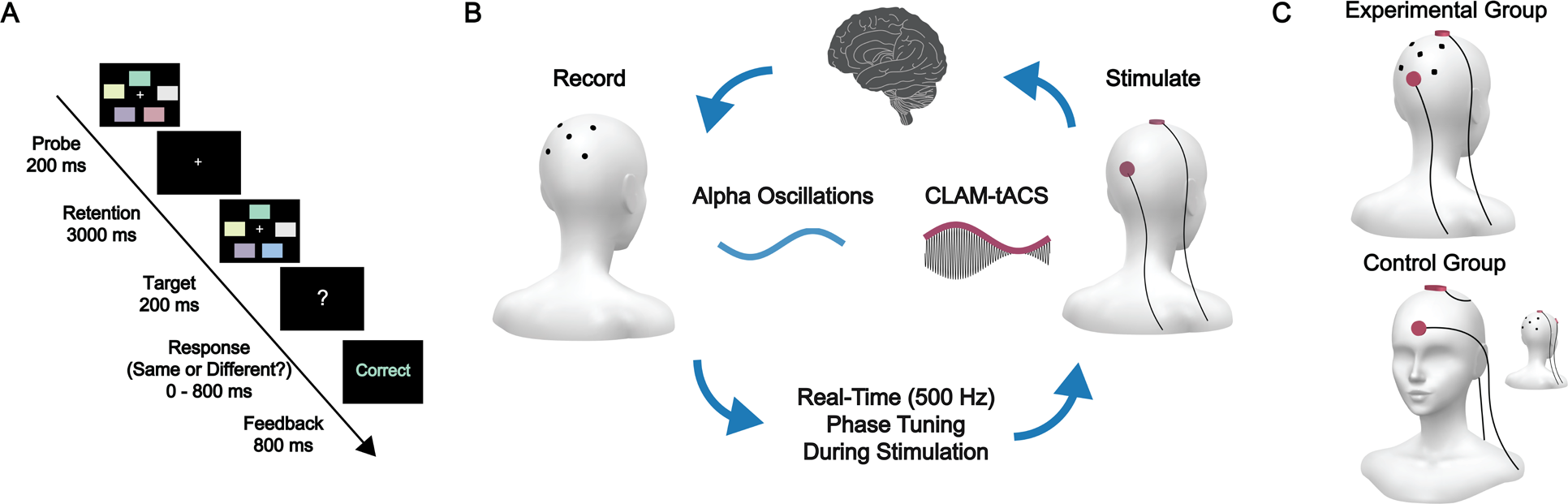
Experimental paradigm. (A) Participants performed a visual WM task (see methods section “Working memory task”) during CLAM-tACS. **(B)** The phase of the envelope of CLAM-tACS (8 kHz carrier frequency and ± 10 mA stimulation amplitude) was adapted to the phase of endogenous alpha oscillations (8 – 14 Hz) in real-time during stimulation, such that their phase lag was maintained around a constant value. For details regarding the real-time signal processing pipeline, see methods section “Real-time EEG processing”. **(C)** In the experimental group, alpha oscillations were recorded from the parietooccipital cortex (Laplacian filter centered on electrode Pz), and CLAM-tACS was applied to the parietooccipital cortex. In the control group, alpha oscillations were recorded from the parietooccipital cortex, but CLAM- tACS was applied to the frontal cortex. The experimental stimulation montage was adapted from prior work [24] because it optimally stimulates sources of parietooccipital alpha oscillations (Fig. S3) known to be linked to visual working memory contents [38, 39].

**Figure 2.**
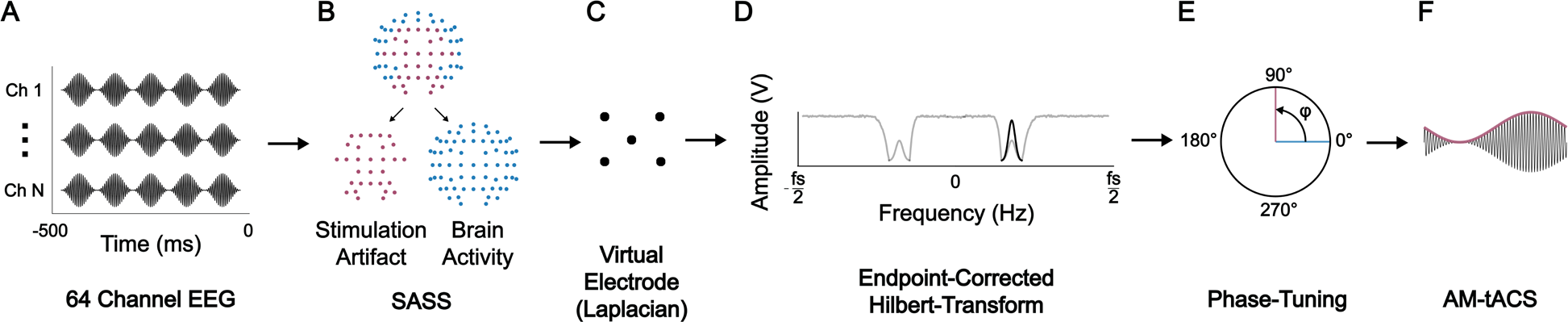
Real-time (500 Hz) signal processing pipeline. **(A)** The latest sample of electroencephalography (EEG) data were appended to a buffer of 500 ms length. **(B)** Stimulation artifact source separation (SASS) was applied to the data to reject artifacts of closed-loop amplitude-modulated transcranial alternating current stimulation (CLAM-tACS). **(C)** A Laplacian (Hjorth) spatial filter centered on the parieto-occipital cortex was computed to obtain a virtual electrode from the data. **(D)** The endpoint-corrected Hilbert transform was applied to the data to remove broadband stimulation artifacts and obtain a real-time phase estimate of parieto-occipital alpha oscillations (8 – 14 Hz). **(E)** The real-time phase estimate of alpha oscillations was incremented by φ to obtain the phase of the CLAM-tACS envelope signal. **(F)** The CLAM- tACS envelope signal was sent to a signal generator, which applied the envelope signal to an 8 kHz carrier signal and sent the resulting waveform to the electric stimulator.

### Participants

Fifty healthy volunteers participated in the experiment, provided they did not exhibit a current or prior neurological disorder, and were randomly assigned to one of the two groups. The study was approved by the ethics committee of the Charité – Universitätsmedizin Berlin (EA1/077/18). Before entering the study, all participants gave written informed consent. Four participants were excluded due to technical reasons (system crashes or insufficient signal quality due to bad electrode contact with the scalp). Data from twenty-one participants (26.1 ± 8.05 years of age, 13 male, 8 female) were included in the evaluation of the experimental group. Data from twenty-five participants (25.4 ± 4.19 years of age, 14 male, 11 female) were included in the evaluation of the control group.

### Electroencephalography

A 64-channel scalp electroencephalogram (EEG) according to the international 10 – 20 system was recorded from each participant in both experiments using a NeurOne system (Bittium Corp., Oulu, Finland). For all recordings, the amplifier was set to DC-mode with a dynamic range of +/-430 mV, a resolution of 51 nV/bit, and a range of 24 bit. Data were sampled at 2 kHz, such that the anti-aliasing low-pass filter had a cut-off frequency of 500 Hz. The EEG system sent all data via real-time UDP protocol to a real time computer, where it was further processed.

### Transcranial alternating current stimulation

Closed-loop amplitude-modulated transcranial alternating current stimulation (CLAM-tACS) was delivered to the scalp with a current of ± 10 mA and a carrier signal frequency of 8 kHz using a Digitimer DS5 (Digitimer Ltd, UK). The envelope signal was phase-tuned to ongoing alpha oscillations in real-time, and thus had a frequency content of 8 – 14 Hz. CLAM-tACS was applied continuously throughout the entire duration of the WM task, and a new phase lag between CLAM-tACS and endogenous alpha oscillations was pseudorandomly chosen out of 6 different phase angles (30°, 90°, 150°, 210°, 270°, and 330°) for each WM trial. The Digitimer DS5 was controlled by a SDG 2042X signal generator (Siglent, NL), which applied amplitude-modulation to the carrier signal depending on an input voltage signal received from the Speedgoat Performance Real-Time Target Machine (Speedgoat GmbH, CH), which executed the real-time signal processing pipeline (Fig. 2). CLAM-tACS was applied through two circular rubber electrodes (34 mm diameter, 2mm thickness) that were attached to the scalp using conductive ten20 paste (Weaver & Co, Aurora, CO, USA). In the experimental group, CLAM-tACS electrodes were centered on positions Oz and Cz of the international 10-20 system. In the control group, CLAM-tACS electrodes were centered on positions Fpz and Cz. Stimulation was neither ramped up nor down. At the end of each session, participants were asked if they perceived any rhythmic skin sensation or visual flicker, which none reported. Since both groups received active stimulation, blinding was not assessed.

### Real-time EEG processing

A Speedgoat Performance Real-Time Target Machine (Speedgoat GmbH, CH) received a real-time UDP stream containing EEG data and executed the signal processing pipeline (Fig. 2) as a Simulink Real-Time R2022a (Mathworks Ltd, USA) multi-rate model. At a rate of 1 Hz, the stimulation artifact source separation (SASS) projection matrix was computed by joint diagonalization of the covariance matrix obtained from data recorded in the presence of CLAM-tACS (past 25 s of continuous stimulation data) and the covariance matrix computed from data recorded in absence of CLAM-tACS (7.5 min before stimulation) [37]. Covariances matrices were computed from narrowband data bandpass-filtered from 8 – 14 Hz using a finite impulse response filter. The following steps were executed at a rate of 500 Hz. First, the latest 500 ms of broadband EEG data were spatially filtered using the current SASS projection matrix. Subsequently, a virtual electrode was computed using a Laplacian (Hjorth) filter [40] with the center electrode Pz and surrounding electrodes PO7, PO8, P3, and P4 according to the international 10-20 system. Finally, the endpoint-corrected Hilbert transform [34] was applied to remove broadband noise and obtain the real-time phase estimate of 8 – 14 Hz target oscillations. The continuous phase-tuning in our approach naturally results in a continuous frequency-tuning, as a constant phase difference between two signals implies that they have identical frequency [41].

### Working memory task

Participants initially performed a visual WM task for 7.5 min in absence of CLAM-tACS (approximately 90 trials). After a 5 min break, they performed the same visual WM task (Fig. 2) for 45 min during CLAM-tACS (approximately 540 trials). Each trial started with the presentation of a white fixation cross in the middle of a black background for 400 ms. Then, five colored squares were displayed on a black background for 200 ms. The squares were positioned on the edges of a regular pentagon, at the center of which a white fixation cross was located. After a retention interval of 3000 ms, a second set of squares was presented for 200 ms. The second set could be identical to the first or differ in color at exactly one location. In half of the trials, a change occurred. Changes in colors and locations were balanced across conditions and groups. Participants had 800 ms to indicate whether the two sets of squares were identical (by pressing j) or different (by pressing f). Subsequently, feedback (“correct” or “incorrect”) was displayed on the screen for 800 ms. Thus, the outcome parameter was whether participants detected the change or not, which was summarized in a single accuracy value per condition.

### Offline EEG processing

MNE-Python [42], NumPy [43], and SciPy [44] were used to evaluate EEG data. First, EEG data were filtered from 8 – 14 Hz using a finite impulse response filter. For EEG data recorded in the presence of CLAM-tACS, stimulation artifact source separation (SASS) was applied to remove the stimulation artifact [45]. Subsequently, a virtual electrode was computed from the Laplacian (Hjorth) filter [40] with the center electrode Pz and surrounding electrodes PO7, PO8, P3, and P4 according to the international 10-20 system. The Hilbert transform was then applied to obtain instantaneous amplitude and phase of alpha oscillations for each trial. For all further analyses, only data from the retention interval was considered.

### Statistics

To mitigate the influence of outliers, we employed Yuen’s t-test for all pairwise comparisons [46], and Shepherd’s method coefficient for all correlations [47].

### Phase-tuning of CLAM-tACS to alpha oscillations

To quantify phase-locking between the CLAM-tACS envelope signal and alpha oscillations, we first computed their phase lag, adjusting for the target phase lag by subtracting it from the actual phase lags. This was done for comparability of phase lags across target phase lag bins. Then, we obtained the maximum likelihood estimate of the concentration parameter (κ) of a von Mises distribution [48] describing the resulting phase lags.

### Modulation amplitude and phase

To assess amplitude and phase of the modulation of WM accuracy and long-range alpha synchrony, the discrete Fourier transform was employed. Let m = [m1, …, mN] represent the mean values of WM accuracy or phase lag index within each phase bin 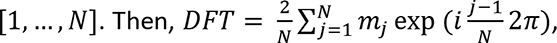, the modulation amplitude |*DFT*|, and the modulation phase ∠*DFT*. To assess correlations between modulation amplitudes, we first normalized them to a percent value, and then computed the Pearson correlation coefficients between them. To assess correlations between modulation phases, we computed the normalized mutual information (NMI) between them [49]. To assess consistency of the lags between modulation phases, we computed the phase locking value (PLV) between them [50]. To assess statistical significance of NMI and PLV, we permuted the modulation phases across participants 1000 times. For each permutation, the test statistic (NMI or PLV) was recomputed. A p-value was then obtained as the fraction of permuted test statistics that were larger than the original test statistic.

### Working memory performance

To evaluate modulation of WM accuracy by CLAM-tACS as a function of phase lag, we used a permutation test [51]. First, for each participant, the WM accuracy was computed for each phase lag between CLAM- tACS and alpha oscillations. Then, WM accuracy was averaged across participants. The discrete Fourier transform was used to obtain a group-level modulation amplitude (see methods section “Modulation amplitude and phase”). To assess statistical significance of this modulation amplitude, we permuted the trials across phase bins within each participant 1000 times. For each permutation, the group-level modulation amplitude was recomputed. A p-value was then obtained as the fraction of permuted modulation amplitudes that were larger than the original modulation amplitude.

### Long-range alpha synchrony

To assess long-range synchronization of alpha oscillations, the phase lag index (PLI) between each pair of channels was computed [52]. To assess modulations of PLI by CLAM-tACS at the network level, network- based statistics were used [53]. Network-based permutation tests follow the same principles as cluster- based permutation tests [54], except that clusters are defined as connected components (networks) in a thresholded graph. First, the modulation amplitude (across CLAM-tACS phase lags) for each pair of channels was computed (see methods section “Modulation amplitude and phase”) to obtain a channel-by-channel square matrix of modulation amplitudes for each participant, which was averaged across participants. Second, the matrix of modulation amplitudes was thresholded at the 99^th^ percentile of its entries. Third, networks (clusters) in this thresholded matrix were identified by finding connected components of the represented graph [55]. To find connected components, the connected components function of the scipy.sparse.csgraph package was employed [44]. A network-level test statistic was then obtained for each network by averaging modulation amplitudes across connections within each network. To assess statistical significance of each network, the single-trial data were permuted across CLAM-tACS phase lag bins 1000 times, and the network-level test statistics (modulation amplitudes) were recomputed. For each permutation, the largest network-level test statistic was saved. A p-value was then obtained for each network as the fraction of permuted network statistics that were larger than the original network statistic.

## Results

### CLAM-tACS was phase-tuned to parietooccipital alpha oscillations

Ofline analysis of EEG data recorded from each participant during CLAM-tACS indicated successful recovery of power spectra (Fig. 3A) and phase information (Fig. 3B to 3D). None of the participants perceived any rhythmic skin sensation or visual flicker. To quantify the performance of real-time phase- tuning, we assessed phase locking between the envelope of CLAM-tACS and parietooccipital alpha oscillations (see methods section “Phase-tuning of CLAM-tACS to alpha oscillations” for a definition of κ) in the target channel, obtained using a Laplacian filter centered on electrode Pz. In absence of electric stimulation, baseline performance of the closed-loop system (simulated with the participant and hardware in the loop) was comparable (t(44) = -0.890, p = 0.378) in the experimental (0.699 ± 0.351 κ) and control (0.780 ± 0.246 κ) groups. During CLAM-tACS, performance of the closed-loop system was lower (t(44) = - 2.68, p = 0.0104) in the experimental (0.411 ± 0.240 κ) compared to the control group (0.584 ± 0.189 κ). Peak alpha frequency in the experimental group (11.1 ± 0.716 Hz) did not differ significantly (t(44) = 1.61, p = 0.12) from that in the control group (10.8 ± 0.869 Hz), indicating that the groups did not differ in the stimulation frequency received.

**Figure 3.**
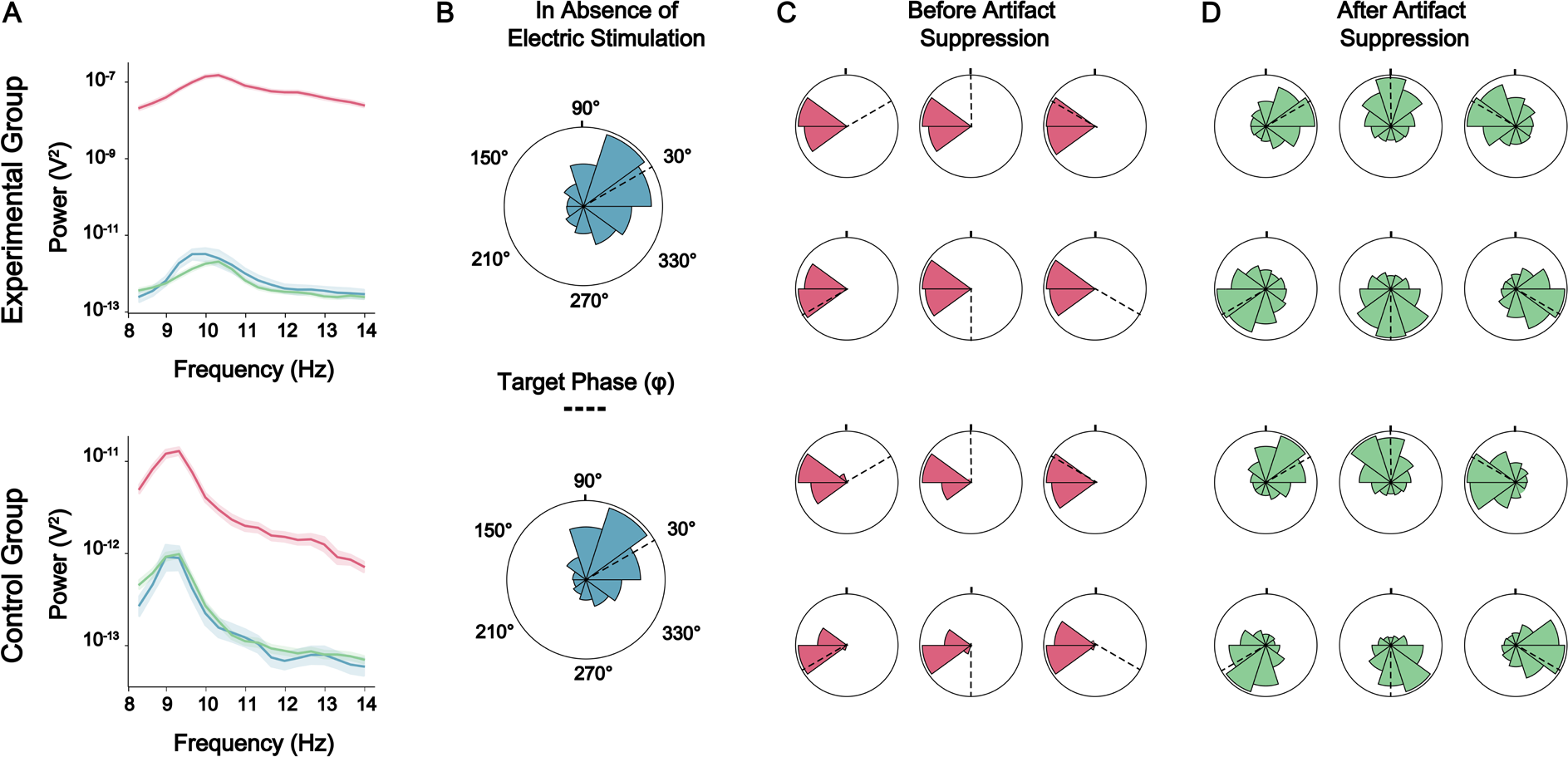
CLAM-tACS was phase-tuned to parietooccipital alpha oscillations in real-time. (**A**) The power spectrum of the target channel in EEG signals recorded during CLAM-tACS was comparable to that in absence of electric stimulation (blue) after application of ofline SASS (green). Shaded areas indicate the 95% confidence interval of the mean. (**B**) In absence of electric stimulation (blue), the closed-loop system tracked the phase of parietooccipital alpha oscillations with the participant and hardware in the loop. (**C**) Before applying SASS offline (red), the phase lag between the envelope of CLAM-tACS and the target channel was locked to approximately 180°, due to the stimulation artifact. (**D**) After applying SASS ofline (green), the phase lag between the envelope of CLAM-tACS and the target channel was locked to the target phase lag. Data of a single participant selected from each group is depicted.

### CLAM-tACS enhanced WM accuracy in a phase lag dependent manner

In the experimental group, we found an effect of the CLAM-tACS phase lag on WM accuracy (p < 0.05, permutation test) (Fig. 4A), while no effect was found in the control group (p = 0.703, permutation test) (Fig. 4B). When assessing the modulation of WM accuracy at a single-participant level using a sine fit across CLAM-tACS phase lags, we found a non-significant trend (t(44) = 0.860, p = 0.198) towards higher modulation in the experimental group (8.39 ± 4.13 %) compared to the control group (7.34 ± 1.82 %). In the experimental group, stimulation with a 330° phase lag to endogenous alpha oscillations most strongly enhanced WM accuracy relative to the pre-tACS baseline (t(20) = 3.38, p_Bonferroni_ = 0.0164, d_z_ = 0.976). Stimulation at the opposite phase lag (150°) did not suppress performance relative to baseline (t(20) = - 0.0736, p_Bonferroni_ > 1, d_z_ = -0.0212). Notably, however, we found that the phase lag resulting in the greatest enhancement of WM accuracy (326 ± 98.3°) varied across participants (Fig. 6B), as identified by fitting a sine function to the data across phase lag bins (see methods section “Modulation amplitude and phase”). Note that the phase lags for enhancement and suppression were identified jointly as the location of the maximum and minimum of this sine function.

**Figure 4.**
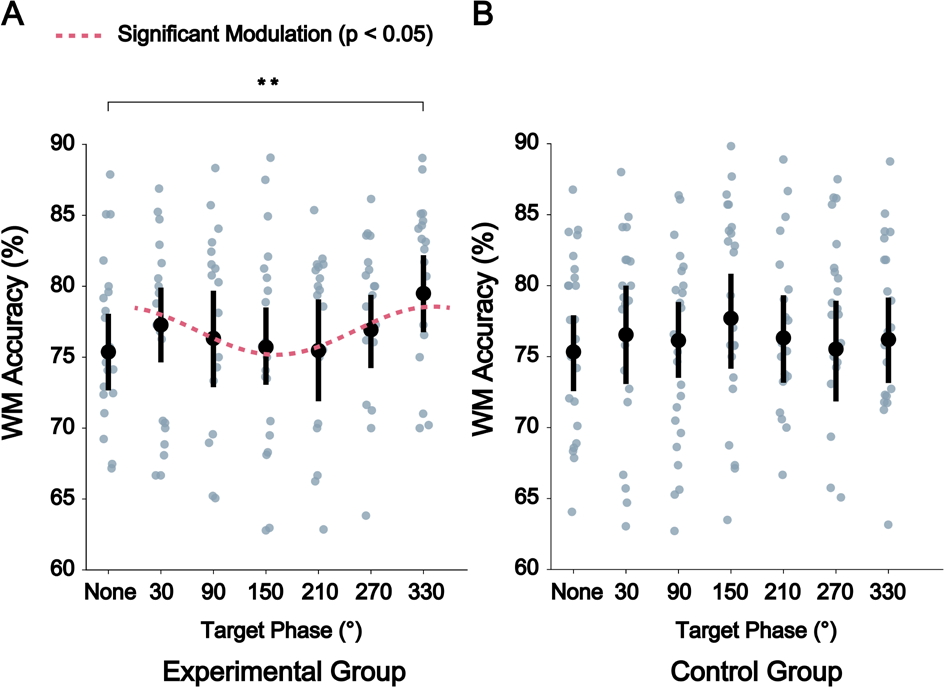
CLAM-tACS enhanced WM accuracy in a phase-dependent manner. (**A**) In the experimental group, WM accuracy was modulated by the CLAM-tACS phase lag (p < 0.05, permutation test). Stimulation with a 330° phase lag to endogenous alpha oscillations most strongly enhanced WM accuracy relative to baseline (t(20) = 3.38, p_Bonferroni_ < 0.05, d_z_ = 0.976). Stimulation at the opposite phase lag (150°) did not suppress performance relative to baseline (t(20) = -0.0736, p_Bonferroni_ > 1, d_z_ = -0.0212). (**B**) In the control group, no effect of CLAM-tACS phase lag on WM accuracy was found (p = 0.703, permutation test).

### CLAM-tACS concurrently modulated frontoparietal alpha synchrony and WM accuracy

We then assessed effects of the CLAM-tACS phase lag on alpha synchrony, as measured in EEG sensor signals. Using network-based statistics (see methods section “Long-range alpha synchrony”), a frontoparietal network of brain regions was found to have been modulated by CLAM-tACS (p < 0.001, permutation test) (Fig. 5A). When assessing the percent modulation of alpha synchrony in this network at a single-participant level using a sine fit across CLAM-tACS phase lags, we found a non-significant trend (t(44) = 1.02, p = 0.159) towards a higher modulation in the experimental group (5.08 ± 1.20 %) compared to the control group (4.51 ± 1.58 %). In the experimental group, the phase lag enhancing alpha synchrony in this network varied across participants (223 ± 114°) (Fig. 5B) and correlated with the phase lag enhancing WM accuracy (r_nmi_ = 0.610, p < 0.05, permutation test). Notably, these phase lags exhibited a consistent (PLV = 0.415, p < 0.05, permutation test) lag (215 ± 76.0 °) (Fig. 5D), indicating that whenever frontoparietal alpha synchrony was enhanced, WM accuracy was suppressed, and vice-versa. Note that the phase lags for enhancement and suppression were identified jointly as the location of the maximum and minimum of a sine function fit to the data across phase bins (see methods section “Modulation amplitude and phase”). Importantly, the amplitude of alpha synchrony modulation also correlated with the amplitude of WM accuracy modulation (r = 0.456, p < 0.05) (Fig. 5C). In the control group, no modulation of alpha synchrony by CLAM-tACS was found (p = 0.130, permutation test).

**Figure 5.**
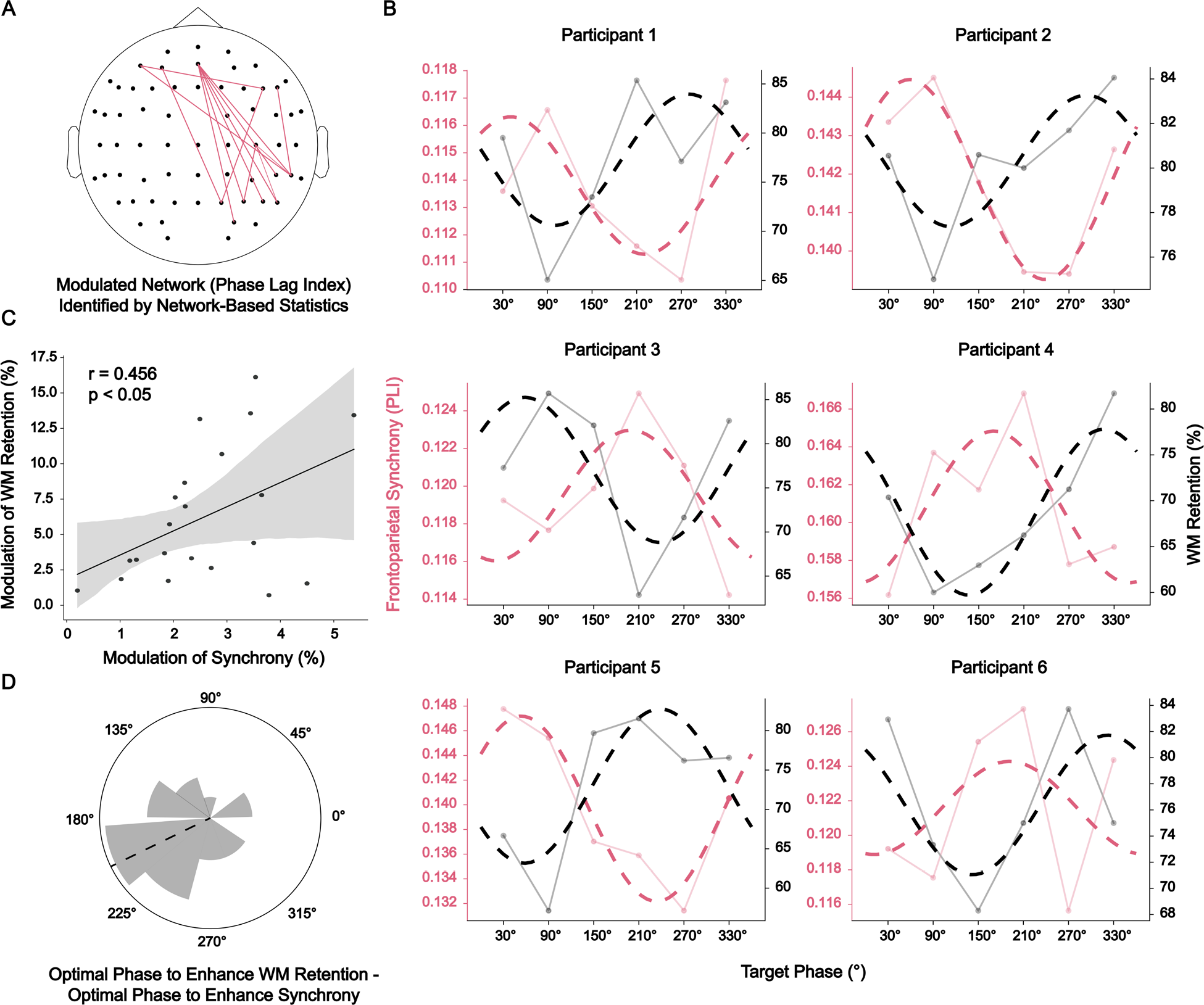
CLAM-tACS concurrently modulated frontoparietal alpha synchrony and WM accuracy. (A) Long-range synchrony of alpha oscillations in a frontoparietal network of brain regions was modulated by the CLAM-tACS phase lag (p < 0.001, permutation test). **(B)** The phase lags that most strongly enhanced frontoparietal alpha synchrony (red) and WM accuracy (black) varied across participants. For analysis of modulation amplitude and phase, a sinusoidal function (dotted line) was fit to the data (see methods section “Modulation amplitude and phase”) averaged over trials within each participant. **(C)** The modulation amplitude of frontoparietal alpha synchrony predicted the modulation amplitude of WM accuracy (r = 0.456, p < 0.05). Each data point represents one participant. **(D)** The phase lag leading to enhancement of frontoparietal alpha synchrony predicted the phase lag leading to enhancement of WM accuracy (r_nmi_ = 0.610, p < 0.05, permutation test). These phase lags exhibited a consistent (PLV = 0.415, p < 0.05, permutation test) lag (215 ± 76.0°).

## Discussion

### CLAM-tACS enhances working memory in healthy adults

Our results demonstrate that continuous phase-tuning of CLAM-tACS is feasible and can effectively enhance WM accuracy in healthy adults. Our results are consistent with earlier work, where in-phase (0°) stimulation of parietooccipital alpha oscillations led to enhancement of WM accuracy when compared to anti-phase (180°) stimulation [13]. In comparison to this earlier study with a small to medium effect size (d_z_ = 0.354), a large effect size of d_z_ = 0.976 was achieved using CLAM-tACS. This suggests that continuous phase-tuning of CLAM-tACS can reduce behavioral effect variability. However, due to differences in experimental parameters between our study and prior work, this hypothesis must be tested in a future comparative experiment. Further studies are also required to assess any potential benefit of CLAM-tACS compared to other tACS protocols that either individualize the stimulation frequency [16] or combine tACS with TMS [56] to enhance WM.

### CLAM-tACS enables phase-dependent enhancement and suppression of brain oscillations

The phase lag between tACS and endogenous brain oscillations is increasingly recognized as an important mediator of stimulation effects [22, 29, 32–34, 57]. Like prior work, we found that effects of tACS on WM accuracy are linked to phase lag dependent enhancement and suppression of frontoparietal alpha oscillations. While tACS is typically frequency-tuned to endogenous brain oscillations by recording EEG/MEG before stimulation [11, 58–62], continuous phase-tuning of tACS has been impeded by stimulation artifacts [28, 29, 37]. A notable exception is the suppression of tremor by motor cortex or cerebellar tACS that was achieved by phase-tuning tACS to the peripherally recorded motor signals [33, 34], which provided an indirect read-out of oscillatory brain activity. In contrast, CLAM-tACS facilitates phase-dependent enhancement and suppression of brain oscillations recorded directly from the scalp, paving the way for novel scientific and clinical applications, particularly those where suppression of pathological oscillations is desired. The ability of CLAM-tACS to suppress pathological oscillations should be investigated in future studies. It is also notable that the above prior studies on reduction of tremor by tACS phase-tuned to peripheral signals reported plastic changes that occurred in the stimulated circuits over the course of stimulation, as evidenced by tremor suppression sustained beyond the stimulation period [34]. It is therefore conceivable that CLAM-tACS could lead to a potentiation of the frequently reported tACS-aftereffects on the target oscillation [35, 63, 64]. This could be examined in future studies and would require applying CLAM-tACS at a single phase lag for a prolonged period, as opposed to the trial-by-trial pseudorandomization of phase lags that was implemented here.

### Suppression of frontoparietal alpha synchrony during CLAM-tACS is linked to improved WM accuracy

We found that enhanced WM accuracy during CLAM-tACS was accompanied by suppressed frontoparietal alpha synchrony during CLAM-tACS (Fig. 5). Thus, our results suggest that WM accuracy is linked to suppression of frontoparietal alpha synchrony, possibly by releasing task-relevant networks from inhibition [65–67]. This supports the theory that alpha oscillations play an inhibitory role in sensory information processing [12, 66, 68], rather than actively maintaining information in WM [8, 69–71], and highlight the importance of assessing brain signals during tACS to understand the causal link between brain oscillations and behavior [28, 72]. Please also note that our task did not include to-be-suppressed irrelevant WM items (Fig. 2), which may explain why suppression of oscillatory activity in the alpha band may be beneficial in our task context (as opposed to WM tasks with distractors [12]). Importantly, we found that stimulation at the optimal phase lag (330°) enhanced WM accuracy, while stimulation at the opposite phase lag (150°) did not result in impaired WM accuracy (Fig. 4). We speculate that this may relate to the dependence of tACS effects on the strength of synchrony in targeted oscillations, as previously shown [32]. If synchrony in ongoing oscillations is strong, tACS may only be able to suppress the targeted oscillation and associated behavior. Conversely, if synchrony in ongoing oscillations is weak, tACS may only be able to enhance the targeted oscillation. However, this conjecture remains to be tested within a dedicated study with an appropriate experimental design.

### Advantages of CLAM-tACS over conventional tACS

While CLAM-tACS and conventional tACS can both enhance and suppress targeted brain oscillations, their relative efficacy remains to be investigated. When conventional tACS is applied at a fixed frequency that is matched to the endogenous frequency of targeted brain oscillations, it can enhance ongoing activity [23, 24]. In contrast, when conventional tACS is applied at a fixed frequency that is slightly higher or lower than the endogenous frequency of targeted oscillations, it can suppress ongoing activity [73]. In contrast, CLAM- tACS is adapted to brain activity in real-time, enabling phase-dependent enhancement and suppression of targeted brain oscillations [29]. In general, it is unclear whether the frequency-dependent mechanism of conventional tACS or the phase-dependent mechanism of CLAM-tACS is more effective at enhancing or suppressing brain oscillations or modulating brain function and behavior. Only a side-by-side comparison of both protocols under equal experimental conditions will provide an actionable answer to this question. Since the assessment of brain activity recorded during conventional tACS is exceedingly difficult, we propose the assessment of online behavioral effects and immediate physiological aftereffects for such a side-by-side comparison. Sensory stimulation confounds may differ between the stimulation protocols, however, and can further complicate such a comparison.

CLAM-tACS may also enable entirely novel scientific investigations. For instance, it is known that key brain functions, such as the representation of position in space [74], maintenance of WM contents [75], or the sampling of visual inputs by attention [76], are locked to distinct phases of low-frequency brain oscillations. However, it is difficult to obtain evidence for such phase codes non-invasively, as the neuronal correlates of cognition and behavior must be measured in a time-resolved manner. Furthermore, existing investigations are not able to experimentally manipulate such phase codes. By stimulating at distinct phases of a low-frequency brain oscillation using CLAM-tACS, and assessing the resulting behavioral effects, it may be possible to further uncover the nature by which the brain uses oscillations to temporally organize information processing and behavior [77].

## Limitations

Both our experimental design and technical approach warrant further examination. As the control group also received CLAM-tACS phase-locked to parietooccipital alpha oscillations (but stimulating frontal regions), it is possible that a direct (transcranial) modulation of frontoparietal synchrony (and WM accuracy) occurred in the control condition, but never reached significance. Furthermore, although the highest electric field intensity in our experimental stimulation montage was in primary visual areas (Fig. S3), we found a modulation of frontoparietal alpha synchrony. Thus, a stimulation montage specifically targeting the parietal cortex may have been more effective at modulating frontoparietal synchrony in the experimental condition. Therefore, it remains unclear why we find only a non-significant trend towards a stronger modulation of alpha synchrony and WM accuracy in the experimental compared to the control group (see sections “CLAM-tACS enhanced WM accuracy in a phase lag dependent manner” and “CLAM- tACS concurrently modulated frontoparietal alpha synchrony and WM accuracy”). Additional control conditions, e.g. stimulation of the shoulder [78] or transorbital stimulation of the retina [57], are required to fully disentangle the contributions of transcranial and sensory stimulation mechanisms in this paradigm and future CLAM-tACS experiments. Furthermore, an increase in WM accuracy during CLAM-tACS relative to baseline in our study suggests that CLAM-tACS enhanced WM accuracy, but an additional sham stimulation control condition would have provided stronger evidence. Finally, stimulation at exactly 0° phase lag between CLAM-tACS and alpha oscillations would have allowed for a better comparison of our results with prior work on phase-dependent effects of tACS on WM [13].

Although CLAM-tACS offers several advantages over conventional tACS, several key issues remain to be resolved. First, it should be noted that CLAM-tACS does not fully eliminate the problem of residual artifacts. Utmost caution is warranted when interpreting EEG signals recorded in the presence of electric stimulation, even after artifact suppression [79–81]. To minimize the influence of residual artifacts on our estimate of alpha synchrony, we have utilized the phase lag index [52], which ignores 0° and 180° phase differences resulting from interference common to both signals. Most importantly, however, we have obtained a tight link between the modulation of alpha synchrony and the modulation of WM accuracy (Fig. 5). In general, when interpreting EEG data recorded during electric stimulation, we argue that it should be mandatory to corroborate any observed physiological effect by means of a direct (online) behavioral effect. Furthermore, in the version of the system presented here, phase-tuning only reached about 60% of performance in absence of electric stimulation, and performance in the experimental group was worse than in the control group (see results section “CLAM-tACS was phase-tuned to parietooccipital alpha oscillations”). Initial tests indicate that performance of phase-tuning during CLAM-tACS can be improved by using a linearly-constrained minimum-variance beamformer [82] for artifact suppression and source reconstruction (unpublished data), as applied in [83] and [84]. The use of a beamformer to suppress stimulation artifacts has the added benefit of allowing for connectivity estimates in source-space (unlike the sensor-space analyses presented here), which is preferable to mitigate volume conduction [52]. Additionally, in applications where CLAM-tACS should be used purely for enhancement or suppression of brain activity and behavior, a calibration session could be performed to identify the optimal phase lag for each participant, e.g. using neuroadaptive Bayesian optimization [85, 86].

Finally, successful phase-tuning during CLAM-tACS depends on the signal-to-noise ratio (SNR) of the target oscillation [87]. The use of spatial filters designed to maximize SNR [88] may facilitate the application of CLAM-tACS to brain oscillations in other frequency ranges, e.g. in the theta or beta band.

While these technical challenges must be addressed to render CLAM-tACS viable for widespread use, the associated reduction of effect variability may enable novel investigations of brain – behavior relationships, as well as novel clinical applications. Here, it will be important to directly compare the efficacy of CLAM- tACS against established open-loop tACS protocols to quantify its benefits.

## Conclusion

We presented a new method that allows for real-time phase-tuning of AM-tACS to ongoing brain oscillations during stimulation (CLAM-tACS). We validated CLAM-tACS by modulating parietooccipital alpha oscillations during a visual WM paradigm, where the phase lag between CLAM-tACS and endogenous alpha oscillations was systematically varied across trials. We found that the effect of CLAM-tACS on WM accuracy was dependent on the phase lag and was linked to online changes in frontoparietal alpha synchrony. While the technical performance of our approach must yet be fully characterized and potentially improved, real- time phase-tuned tACS holds promise for targeted enhancement and suppression of oscillatory brain activity and associated functions [13, 33, 34].

## Author contributions

**David Haslacher**: Conceptualization, Methodology, Software, Investigation, Writing, Data Curation, Visualization. **Alessia Cavallo**: Conceptualization, Methodology, Writing, Data Curation, Visualization. **Philipp Reber**: Conceptualization, Investigation, Writing. **Anna Kattein**: Investigation, Writing, Visualization. **Moritz Thiele**: Investigation, Writing, Editing. **Khaled Nasr**: Writing, Editing. **Kimia Hashemi**: Investigation. **Rodika Sokoliuk**: Writing, Editing. **Surjo Soekadar**: Conceptualization, Project administration, Funding acquisition, Writing, Editing.

