## Supplementary Materials for "Working memory enhancement using real-time phase-tuned transcranial alternating current stimulation"

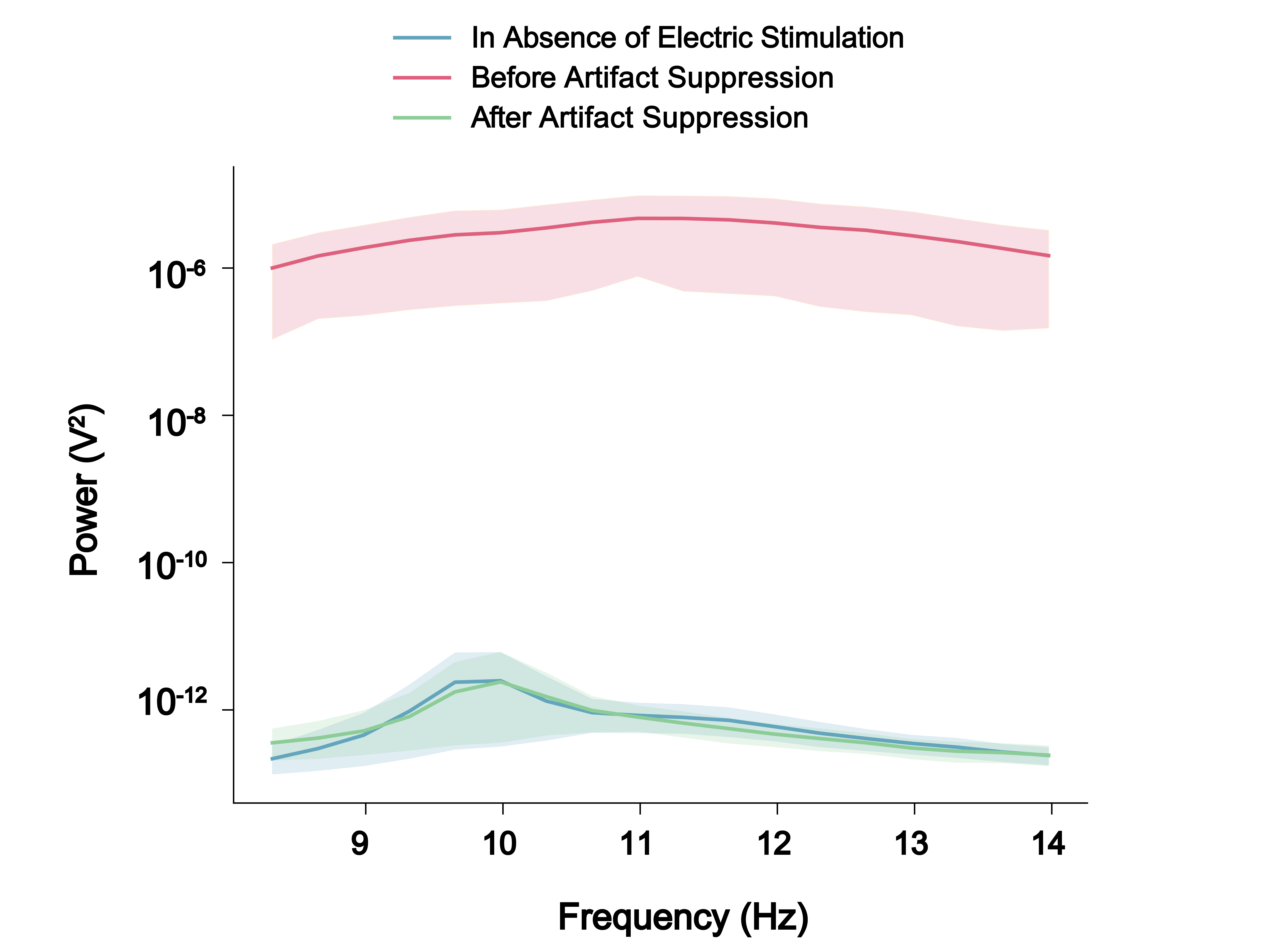

**Figure S1.** Group-averaged power spectrum. In absence of electric stimulation, the group-averaged power spectrum of the target channel (obtained by applying a spatial filter to the EEG data, see Fig. 2) exhibited a peak at approximately 10 Hz. During CLAM-tACS, before artifact suppression, a large artifact was evident in the alpha frequency range. After artifact suppression, the power spectrum of physiological brain activity was recovered. Data from the occipital stimulation condition is depicted.

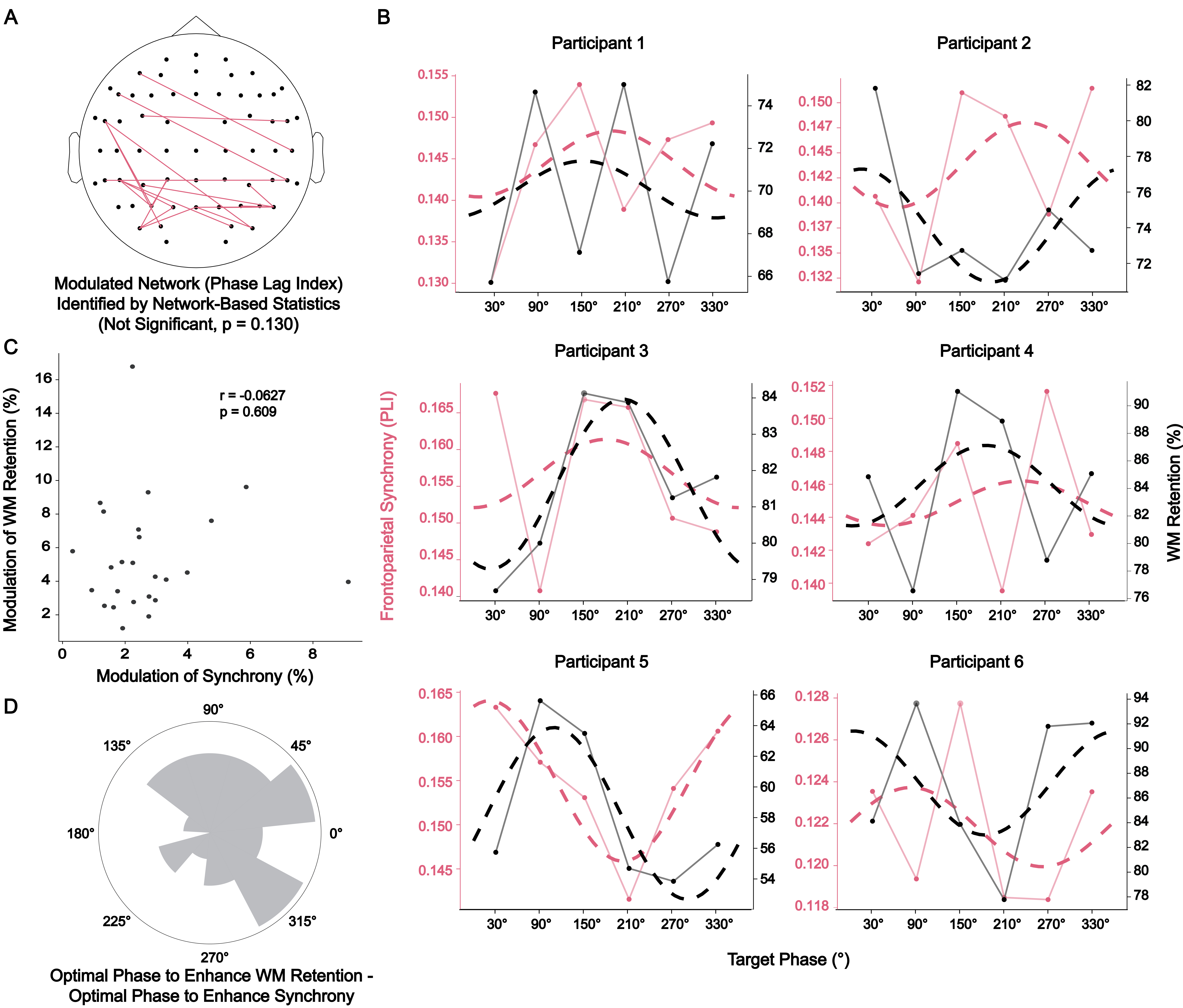

**Figure S2. CLAM-tACS did not concurrently modulate frontoparietal alpha synchrony and WM performance in the control group. (A)** Like WM performance (p = 0.703, permutation test), long-range synchrony of alpha oscillations (p = 0.130, permutation test) was not significantly modulated by the CLAM-tACS phase lag in the control group. **(B)** The phase lags that were associated with enhanced alpha synchrony (red) and WM performance (black) varied across participants. For analysis of modulation amplitude and phase, a sinusoidal function (dotted line) was fit to the data (see methods section “Modulation amplitude and phase”) averaged over trials within each participant. **(C)** The modulation amplitude of frontoparietal alpha synchrony did not predict the modulation amplitude of WM performance (r = -0.0627, p < 0.609). Each data point represents one participant. **(D)** The phase lag leading to enhancement of frontoparietal alpha synchrony did not predict the phase lag leading to enhancement of WM performance (r_mni_ = -0.0442, p = 0.666). These phase lags did not exhibit a consistent difference (PLV = 0.258, p = 0.19, permutation test).

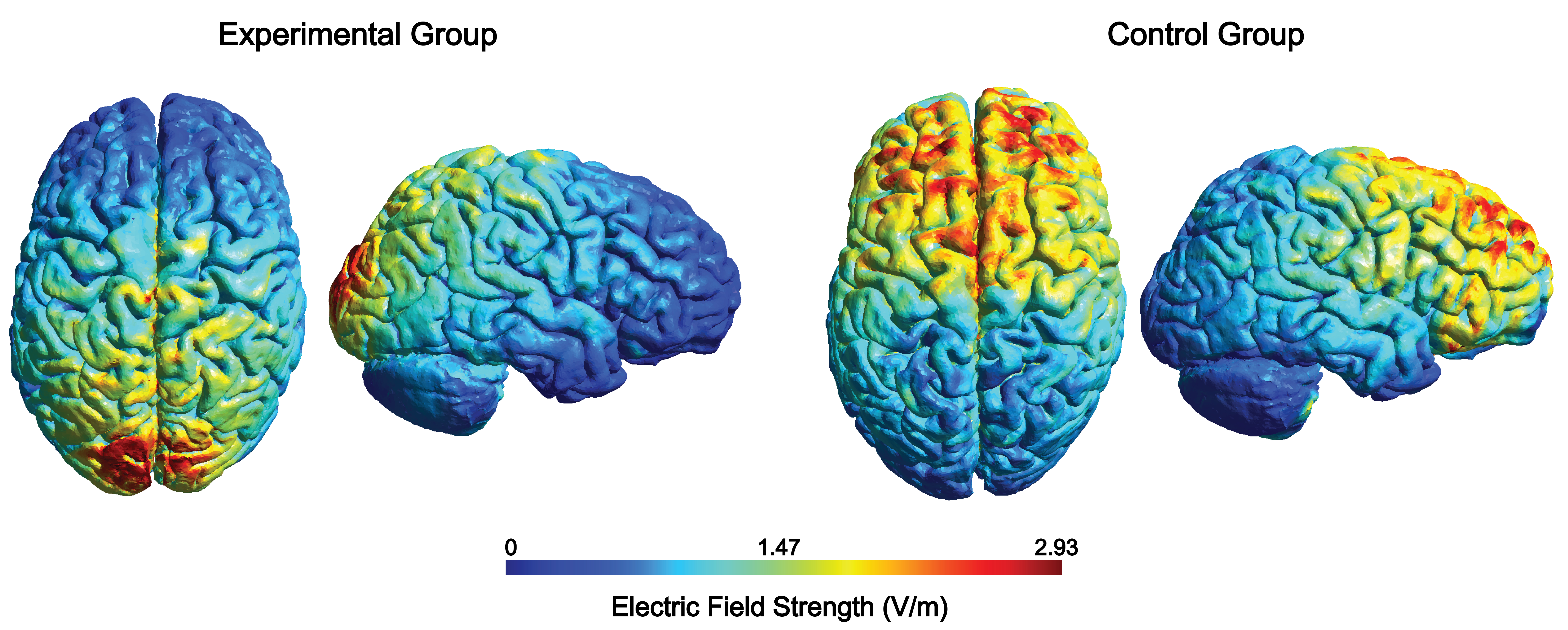

**Figure S3. Electric fields induced by transcranial alternating current stimulation in the experimental and control groups.** In the experimental group, circular rubber electrodes (34 mm diameter) were centered on positions Oz and Cz of the international 10 – 20 system. In the control group, the same electrodes were centered on positions Fpz and Cz. Electric fields were simulated in SimNIBS using a standard head model [1].

[1]	Saturnino GB, Puonti O, Nielsen JD, Antonenko D, Madsen KH, Thielscher A. SimNIBS 2.1: A Comprehensive Pipeline for Individualized Electric Field Modelling for Transcranial Brain Stimulation. In: Makarov S, Horner M, Noetscher G, editors. Brain and Human Body Modeling: Computational Human Modeling at EMBC 2018, Cham (CH); 2019, p. 3-25.
